# Circadian-period variation underlies the local adaptation of photoperiodism in the short-day plant *Lemna aequinoctialis*

**DOI:** 10.1101/2022.03.09.483716

**Authors:** Tomoaki Muranaka, Shogo Ito, Hiroshi Kudoh, Tokitaka Oyama

## Abstract

Phenotypic variation is the basis for trait adaptation via evolutionary selection.^1, 2, 3^ However, the driving forces behind the quantitative trait variations remain unclear owing to their complexity at the molecular level.^4, 5, 6^ This study focused on the natural variation of the free-running period (FRP) of the circadian clock because FRP is a determining factor of the internal clock phase (chronotype), which is responsible for physiological timing during a day.^7^ Although natural variations in FRP have been widely reported,^8, 10, 11^ few studies have shown the association between FRP and adaptive temporal traits. As a clock-dependent physiological process, photoperiodism is a typical target of local adaptation.^12, 13^ *Lemna aequinoctialis* in Japan is a paddy-field duckweed exhibiting a latitudinal cline of critical day-lengths (CDLs) for short-day flowering.^14^ To investigate the relationship between FRP and CDL, we collected 72 strains of *L. aequinoctialis* within a latitudinal range between 31.5°N to 43.8°N. We found a significant correlation (*P* = 7.5E-8) between FRPs and locally adaptive CDLs, confirming that the variation in FRP-dependent chronotypes underlies geographically differentiated photoperiodism. Diel transcriptome analysis revealed that the induction timing of a florigen gene is key for connecting chronotypes to photoperiodism at the molecular level. Based on these results, we propose a fundamental rule concerning the “chronotype effect” in evolution: the variation of FRP functions as a resource for the variation of temporal traits. This study highlights the adaptive significance of FRP variation and provides a reason for the maintenance of FRP variation in natural populations.

## Results and discussions

We investigated the influence of natural variations in circadian clock properties on the temporal phenotype of clock-dependent physiology. The circadian clock is an endogenous timing system that generates a self-sustained daily oscillation entrained to day-night cycles via environmental cues.^15^ The eukaryotic circadian clock consists of feedback loops of multiple clock genes.^16^ The free-running period (FRP) of the circadian clock is a polygenic trait that exhibits natural variation.^8, 9, 10, 11, 12, 17^ These circadian clocks with different periods are entrained to 24 h day-night cycles.^18, 19^ The peak timing (phase) of an entrained rhythm also shows natural variation. This phase phenotype is termed the chronotype.^7^ In this study, we define the “chronotype effect” as the phenotypic effects on physiological traits originating from variations in chronotype. Previous studies suggest that chronotype determination is related to the FRP. ^20, 21, 22, 23, 24, 25, 26^ We hypothesized that FRP variation is associated with local adaptation of a clock-dependent physiology.

*Lemna aequinoctialis* in Japan is a paddy-field duckweed with a latitudinal cline of critical day-lengths (CDLs) for short-day flowering.^14^ To investigate the relationship between FRP and CDL, we collected 72 strains of *L. aequinoctialis* from 20 populations located between latitudes 31.5°N and 43.8°N (Figure 1A and Table S1). To estimate both intra-population and inter-population variation, 11–12 strains were collected from each of four populations (N32Ka, N35Ht, N35So, and N44Ha). The strains were aseptically maintained by clonal growth. Laboratory experiments using a clonal plant enabled the precise determination of multiple phenotypic traits of the natural genotype. The frond size showed intra-population and inter-population variation, suggesting heterogeneous genotypes, even within a population (Figure S1). The estimated genome sizes of all the strains were similar (Figure S2).

**Figure 1.**
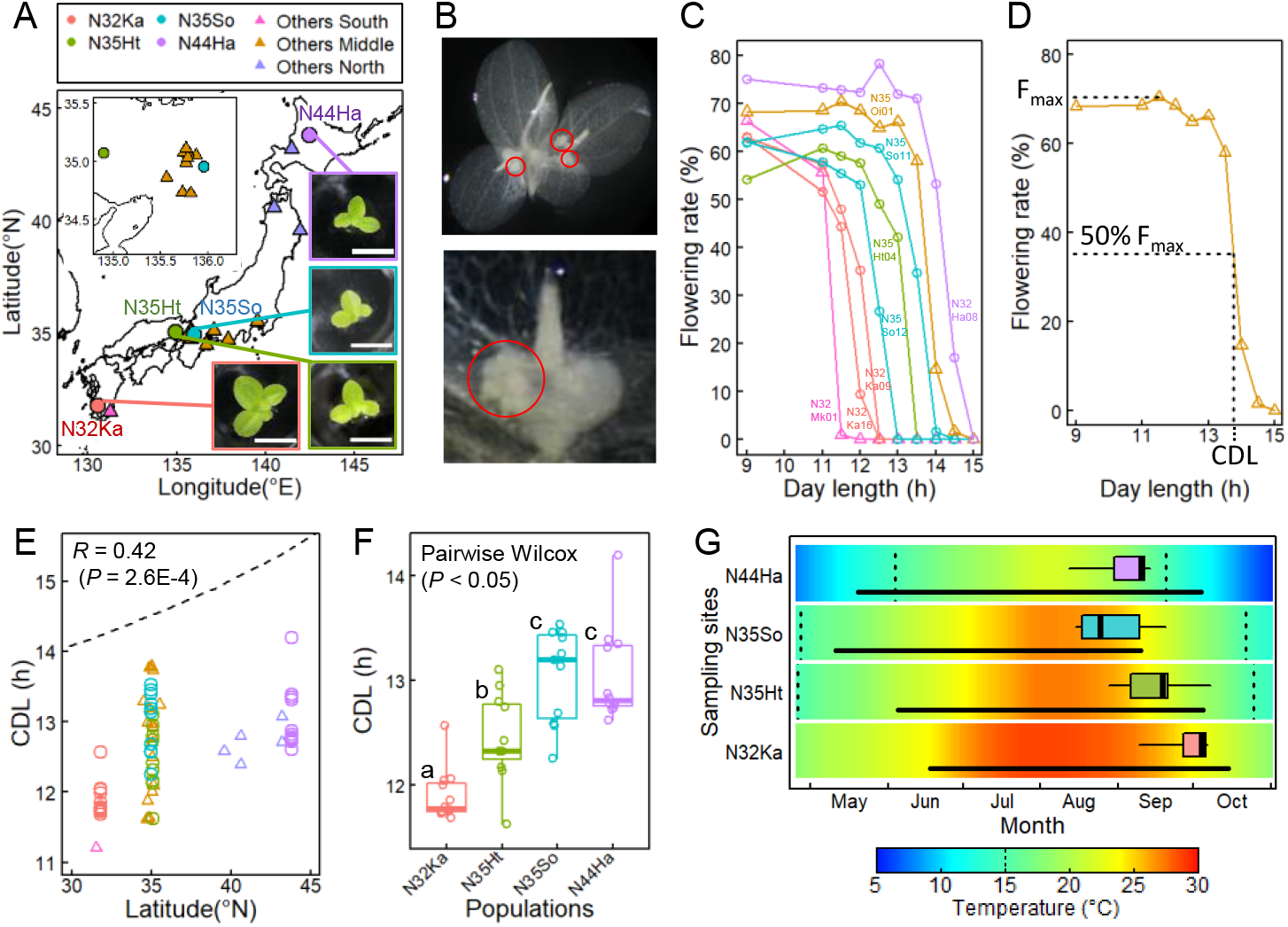
Local adaptation of critical day-length (CDL) in *Lemna aequinoctialis* strains isolated in Japan. **A**, The locations from which 11–12 strains (circles) and 1–3 strains (triangles) were isolated. The same colors and symbols apply to other panels. Examples of the strains in the four populations are shown. Scale bars: 5 mm. **B**, A colony with floral buds (top). Colony of N35Oi03 strain bleached by EtOH after one week of short-day (9L15D) treatment and imaged from the underside. Close-up underside view around the meristematic tissue. Red circles indicate floral buds. **C,** Continuous variation of photoperiodic response on flowering. Flowering rate after one week of photoperiodic treatment is plotted against treated day-length. Eight strains in six populations are shown. **D**, Critical day length (CDL) was determined as the day-length where 50% of the maximum flowering rate (Fmax) is expected. See Methods for more detail. **E**, Latitudinal cline of CDL. The Pearson’s correlation coefficient and *P* value are shown. Dashed line represents day-length of summer solstice. **F,** CDL variation among the four populations. Boxplots with points representing individual strains are shown. **G,** Timing of CDLs in the year at four sampling sites. The dates for CDLs shown in panel c are represented as a box plot at each site. Black horizontal lines represent the flooding period. The background colors represent 30-year (1990–2019) mean of daily temperature. Vertical dotted lines represent 15 °C.

These strains recognized day lengths with a resolution of 0.5 h and their CDLs ranged from 11.2 to 14.2 h (Figures 1B, 1C and 1D). As previously reported,^14^ a latitudinal cline was observed in CDLs: 11.2–12.6 h in the south, 12.4–14.2 h in the north, and a large variation of CDLs was observed at approximately 35°N (Figure 1E). The longer CDL at higher latitudes is consistent with other short-day flowering species.^27, 28, 29^ Intra-population differences of 1–1.5 h in CDLs were observed in each population of N32Ka, N35Ht, N35So, and N44Ha (Figure 1F). The significant differences between these populations suggest geographical differentiation of CDLs. The difference in CDL between the two 35°N populations, N35Ht and N35So suggests that the adaptive CDL at each site may differ even at the same latitude (Figure 1F). The flooding period of the paddy fields differed between the sampling sites depending on the cultivation schedule of the different rice cultivars. The longer CDL of N35So, corresponding to earlier flowering, is consistent with the earlier cessation of flooding at this site than that at the N35Ht site (Figure 1G). The CDL of the N32Ka population was shorter than that of the N35 population (Figure 1F). The CDLs of the three populations appeared to be linked to the cultivation schedule at each site (Figure 1G). Complete drainage of paddy fields during and after rice harvesting is a potential selection pressure for duckweed.^30, 31^ Although the rice-harvesting time at the N44Ha site was comparable to that at the N35Ht site, the CDL of N44Ha was longer than that of N35Ht. At the N44Ha site, the low-temperature season (<15 °C: lower limit for *L. aequinoctialis* growth)^32^ began before the rice harvest time, suggesting that low temperature, rather than rice harvest time, may be a potential selection pressure. Overall, the CDLs of the four populations were likely to fit a suitable season for reproduction at each site.

The natural variation in circadian rhythms was assessed using a luciferase reporter assay.^33^ Using particle bombardment, a luminescent reporter, *AtCCA1::LUC*^34^ was semi-transiently introduced into plants grown under long-day conditions (15L9D; 15-h light / 9-h dark). The plants were placed on an automatic luminescent monitoring system and released into constant light (LL) after two days of 15L9D (Figure 2A). All strains showed clear diel luminescence rhythms with morning peaks at 15L9D. Among the four populations, the peak time of N32Ka was significantly later than that of the other three populations (Figure 2B). To estimate FRP under LL, fast Fourier transform-nonlinear least squares (FFT-NLLS) analysis^35^ was used (Figure 2C), in which the rhythm significance was estimated by a relative amplitude error (RAE) that increased from 0 to 1 as the rhythm neared statistical insignificance. Seven strains with a high RAE value (>0.1) or a high standard deviation (SD) of FRP (>1.5 h) were excluded from the following analysis (Figure S3). The FRP of N32Ka was significantly longer than that of the other populations, corresponding to its late peak time (Figures 2B and 2D). This is consistent with the concept of chronotype: a positive correlation between FRP and peak time.^24^

**Figure 2.**
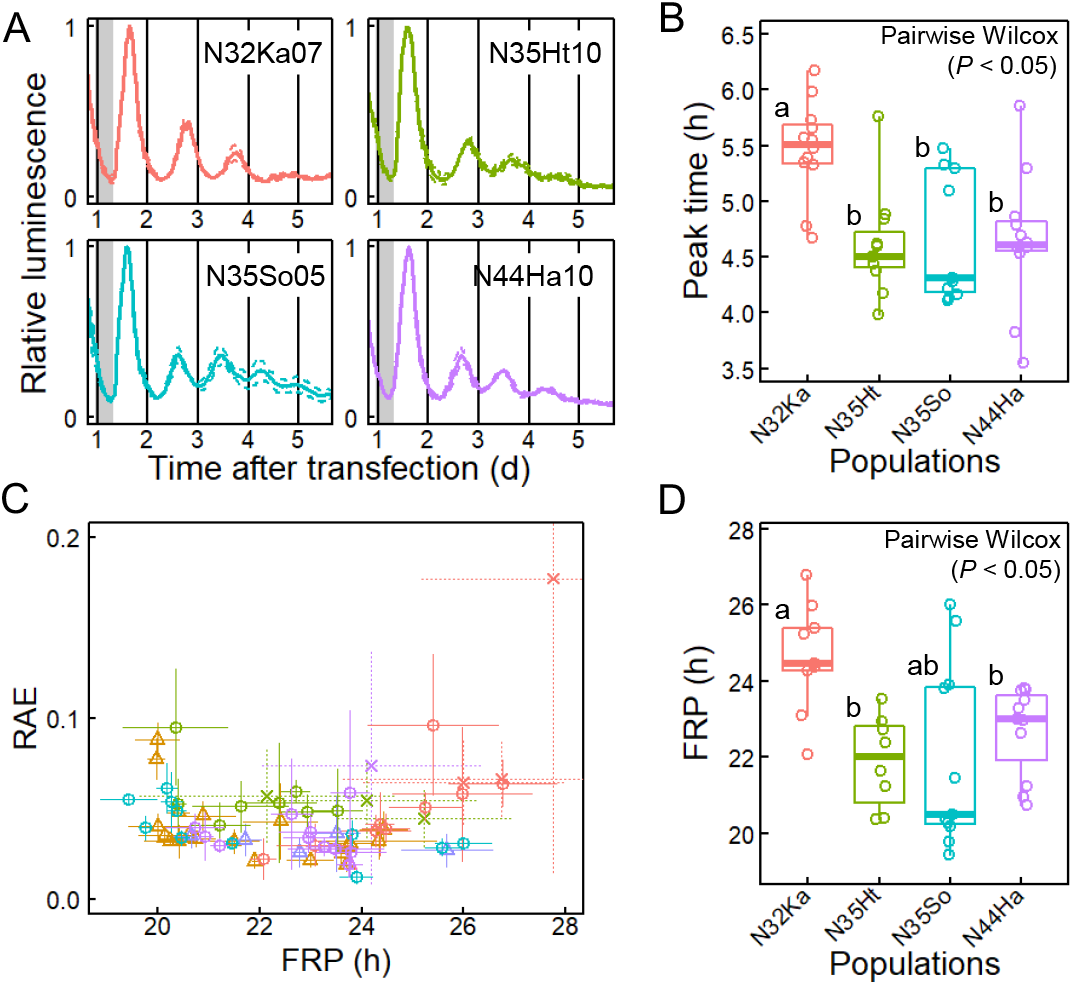
Natural variation of circadian rhythms in *Lemna aequinoctialis* strains. **A**, Examples of luminescence rhythms of *AtCCA1::LUC.* Solid and dashed lines represent mean and ± SD of three replications, respectively. Gray boxes indicate dark period. The strain name is indicated in each plot. **B,** Variation of the timing of the first peak in constant light (LL) among the four populations. Boxplots with points for individual strains are shown. **C**, Relative amplitude error (RAE) of each strain plotted against its free running period (FRP). Mean values of three replications are shown. Error bars = SD. The crosses represent seven strains that showed unstable rhythms (SD of FRP > 1.5 h, Figure S3). Population names are represented in the graph legend. **D,** Variation of the free-running period (FRP) in LL among the four populations. Boxplots with points for individual strains are shown. Seven strains with unstable rhythms (SD of FRP > 1.5 h) were excluded from the plot.

Correlation analysis using data from all 72 strains (20 populations) suggested a chronotype effect on photoperiodism (Figure 3A). A positive correlation between FRP and peak time was confirmed at the genotypic level. In addition, CDL showed a negative correlation with both FRP and the peak time. These results suggest that the variation in FRP-dependent chronotypes underlies the variation in the CDL. The CDL shift can be explained as the chronotype effect on the florigen (floral inducer), which responds to night-length (Figure 3B).^36^ In this model, the activation of the florigen is assumed to be gated by the circadian clock and permitted in the dark. A longer FRP causes a phase delay in the gate timing and, consequently, a shorter CDL, resulting in a negative correlation between FRP and CDL.^37^

**Figure 3.**
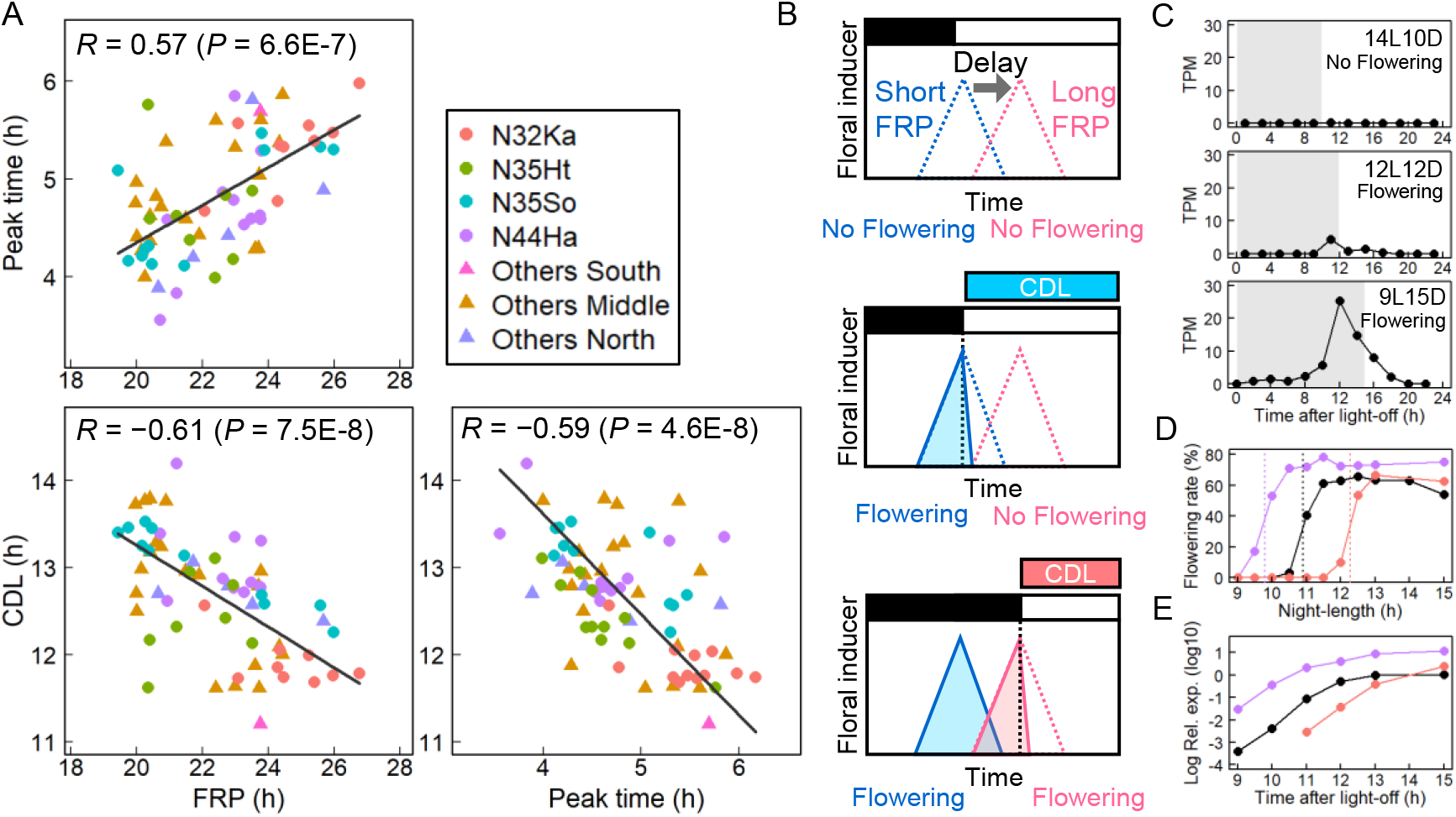
Chronotype effect on photoperiodic flowering. **A,** The correlations between FRP, peak time, and CDL of *L. aequinoctialis* strains. The Pearson’s correlation coefficient and *P* value are shown in each graph. Black lines represent Deming regression lines. Plots of seven strains with unstable rhythms (SD of FRP > 1.5 h) are excluded from graphs that include FRP. **B,** The hypothetical mechanism for the negative correlation between FRP and CDL. The activation of the floral inducer is assumed to be gated by the circadian clock and permitted in the dark. The dotted lines and filled triangles schematically represent gate timing and activity of a floral inducer, respectively. The blue and red colors correspond to strains with a short and long FRP, respectively. **C,** A photoperiodic response of the *LaFTh1* expression. Means of four or two RNA-seq experiments are plotted. Gray box indicates a dark period. **D,** Photoperiodic responses of flowering for three strains (purple, N44Ha08; black, Nd; red, N32Ka06). The flowering rates are plotted against the night length of each photoperiod. The critical night length (24 – CDL) of each strain is shown as dashed line. **E,** *LaFTh1* induction in the three strains during constant dark following 15L9D. The mRNA accumulation quantified by qPCR is plotted. The mRNA expression in N32Ka06 at 9 and 10 h were undetected. Colors are the same as in panel d.

To explore florigen genes, we obtained diel transcriptomes of *L. aequnictialis* pure line (Nd strain^33^) under three photoperiod conditions. Five florigen homologues were detected by *de novo* assembly and were named *LaFTh1–5* (Figures S4A and S4B). *LaFTh1* was induced before the end of the night, and its expression level increased during longer-night conditions (Figures 3C and S4C). The transgenic *Arabidopsis thaliana* carrying an *LaFTh1* overexpression construct showed an early flowering phenotype, suggesting its florigen activity (Figure S4D). *LaFTh1* was induced even in extended dark following a non-flowering long-day condition, suggesting that *LaFTh1* expression directly responds to night lengthening. The timing of *LaFTh1* induction differed among the three strains with different CDLs and the variation in induction timing appears to be responsible for the CDL variation (Figures 3D and 3E). Thus, the induction timing of *LaFTh1* is key for connecting chronotype to photoperiodism at the molecular level. Notably, the *LaFTh1* expression levels showed a large difference. The *LaFTh1* expression in the N44Ha08 strain was 10-fold higher than that in the other two strains (Figure 3E). This high basal level may have caused an extremely long CDL of 14.2 h (Figure 1F).

Our study demonstrated the chronotype effect on locally adapted traits: CDL variation is tightly coupled with FRP variation, even in natural populations. Diel transcriptome analysis suggests that florigen induction timing, which is likely to be related to the FRP-dependent chronotype, is a determining factor for CDL and consequently for the flowering season. These results indicate the importance of circadian phenotypes in the natural selection of seasonal phenotypes. This study also provides new insights into the local adaptation of CDLs to the paddy-field environment. Consistent with previous studies using short-day plants,^27, 28, 29^ the populations of *L. aequnictialis* in Japan showed a latitudinal cline in their CDLs, suggesting that the flowering season of this plant adapted to the local climate, exhibiting a latitudinal cline in temperature.^12^ Furthermore, their CDLs appeared to adapt to the flooding season at each sampling site (Figure 1G). The improved drainage systems associated with the agricultural modernization of Japan that began in the 1950s have resulted in dry paddy-field environments in winter and consequently altered the weed flora.^38^ The natural variation of the FRPs possibly contributed to the adaptive evolution of the flowering season in response to the rapid artificial shift of the flooding season. In addition to inter-population variations, CDLs also showed intra-population variations (Figure 1F). The flooding period of paddy fields is highly dependent on the rice cultivar grown. Such artificial fluctuations may promote intra-population variations in CDLs. In *A. thaliana*, the recombinant inbred lines of two accessions with similar FPRs showed variation as substantial as that observed among the global collection (22.0 to 28.5 h).^39^ Such transgressive segregation of FPR is caused by the complex interference among many quantitative loci,^40^ and may play an important role in providing the phenotypic variations for rapid evolutionary adaptation to irregular environmental changes.^41^ Interestingly, the estimated genome sizes of all *L. aequinoctialis* strains used in this study were ~800 Mbp and considered to be tetraploid (Figure S2).^42, 43^ Polyploidy may generate phenotypic variations by increasing the rates of evolution and number of components in gene regulatory networks.^44, 45^

Because the circadian clock regulates various physiologies,^7, 46^ FRP variation potentially contributes to the fine-tuning of their temporal traits in the process of local adaptation. Conversely, selection pressure on the temporal traits of such clock-dependent physiologies may favor mutations in clock-related genes that alter the FRP and, consequently, the chronotype (Figure 4). This is a possible reason for the maintenance of FRP variation with standing genetic variations in clock-related genes in natural populations.^8, 11, 12^ Thus, the adaptive significance of FRP variation should be considered in relation to the phase phenotype under day-night conditions.^7, 21, 25^ Interestingly, a latitudinal cline of FRP was reported in the linden bug, *Pyrrhocoris apterus*,^10^ in which circadian clock genes are responsible for the photoperiodic response of the reproductive diapause.^47^ The circadian clock was acquired for adaptation to day-night cycles and has evolved as a hub of environmental response systems involving multiple input pathways.^48^ As a result, in the process of adaptive evolution, the circadian system can function as a source of variation for temporal traits through the chronotype effect integrating many quantitative loci for FRP.^4, 39, 49, 50^

**Figure 4.**
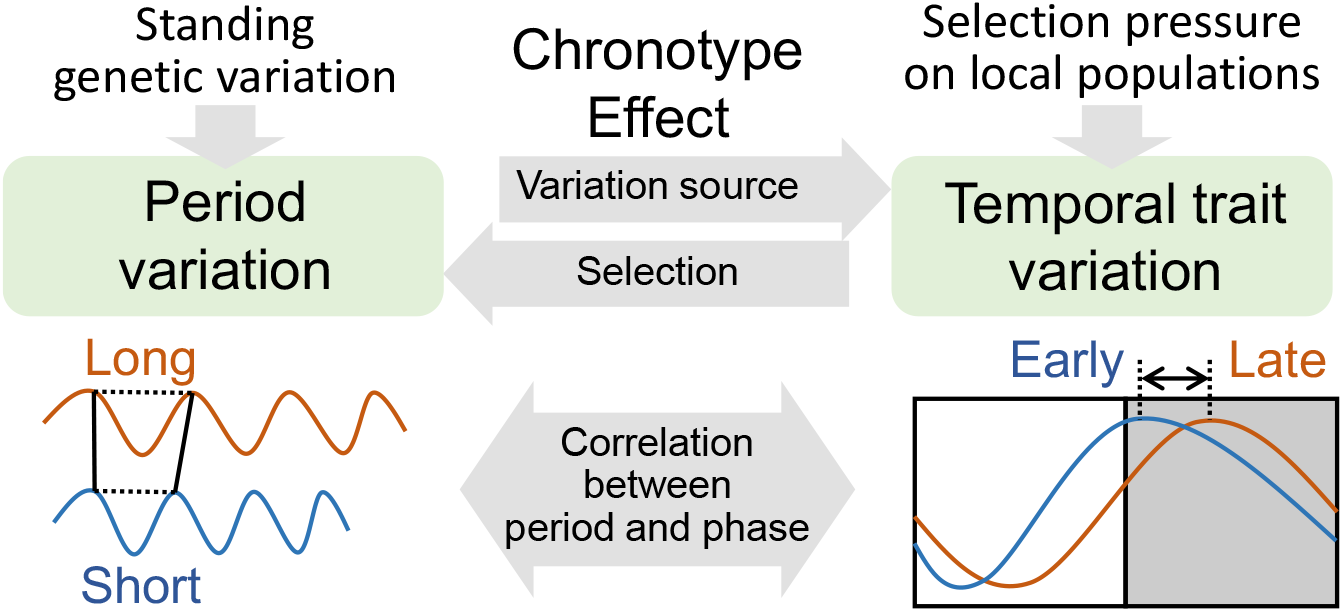
The chronotype effect on local adaptation of clock-dependent temporal traits. FRP variation functions as a resource for the variation of temporal traits in the process of local adaptation. Genetic variations in many quantitative loci for FRP are selected based on their phenotypic effect on temporal traits under day-night conditions. In *L. aequinoctialis*, the selection pressure for early-flowering phenotype appeared to select a short-FRP genotype related to its early induction timing of a florigen gene.

## Data availability

The RNA-seq raw sequences (PRJDB12719) and LaFTh1 CDS (LC662606) were deposited at DDBJ.

## Acknowledgements

We thank N. Emura for her support in collecting duckweed strains, I. Shibano for her support in maintaining duckweed strains, M. Honjo and M. Hirata for their support with RNA-seq experiments in the Joint Usage for Ecological Research, Kyoto University, and S. Kubota for her efforts to establish the Nd strain. This research was supported by the Japan Society for the Promotion of Science KAKENHI [grant numbers JP16H06864 (T.M.), JP20K15861 (T.M.), JP20J00255 (T.M.), JP20K06342 (S.I.), JP21H04977 (H.K), and 17KT0022 (T.O.)], Japan Science and Technology Agency (JST) CREST (JPMJCR15O1, H.K.), JST ALCA (JPMJAL1108, T.O.), and JST SATREPS (JPMJSA2004, T.M., S.I., T.O.).

## Author contributions

T.M. and T.O. designed the study and wrote and revised the manuscript. T.M. performed the experiments using *L. aequinoctialis* and all the analyses. S.I. performed the experiments using *A. thaliana* and contributed to the particle bombardment experiments. H.K. contributed to RNA-seq experiments and revised the manuscript. All authors have contributed to the text.

---

**Figure S1.**
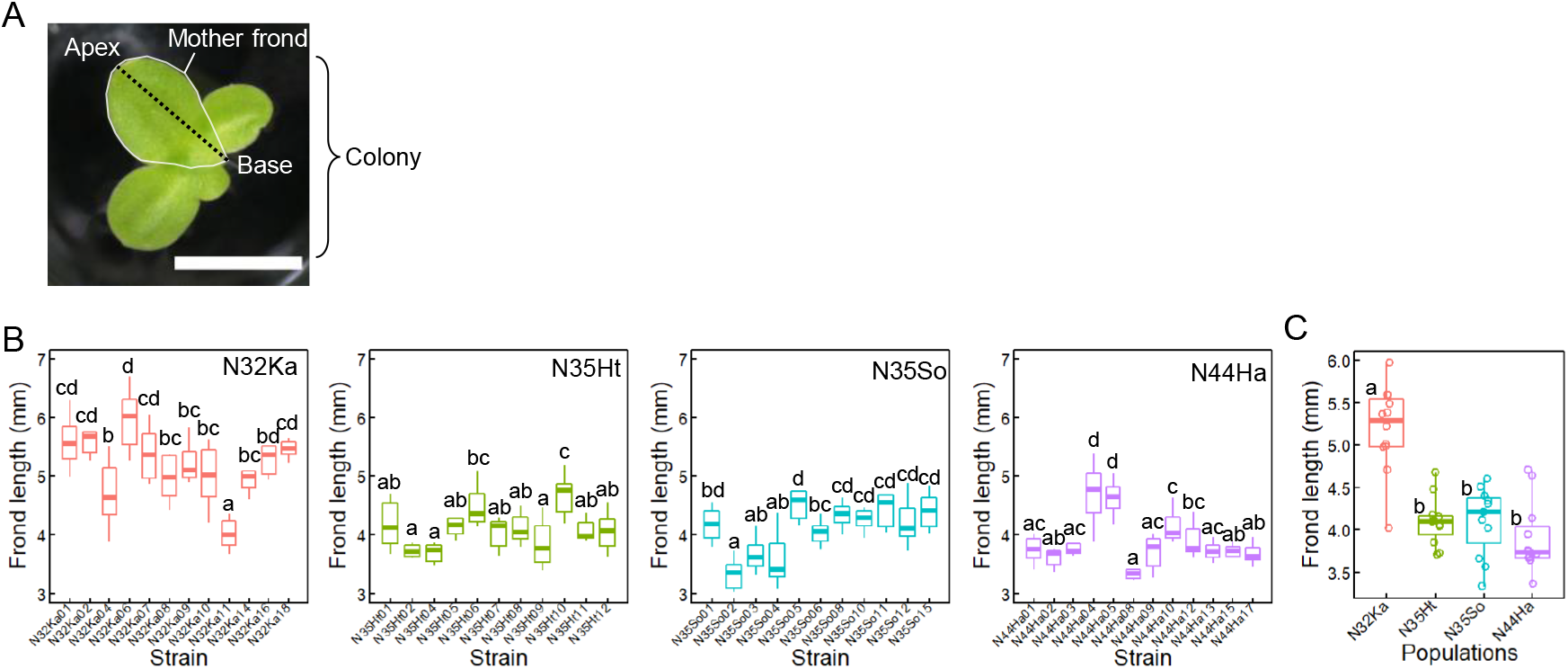
Intra- and inter-population variation of frond length in *L. aequinoctialis* strains. **a,** Definition of a frond length. The length of the mother frond (from base to apex, dotted black line) of eight colonies were measured for each strain of four populations (N32Ka, N35Ht, N35So, and N44Ha). Scale bar indicates 5 mm. **b,** Intra-population variation of frond length of the four populations are shown as boxplots. Different letters indicate significant differences based on Tukey’s HSD test (*P* < 0.05). **c,** Inter-population variation of frond length among the four populations. Different letters indicate significant differences based on pairwise Wilcoxon test (*P* < 0.05). Boxplots display a median line, interquartile range boxes, min/max whiskers.

**Figure S2.**
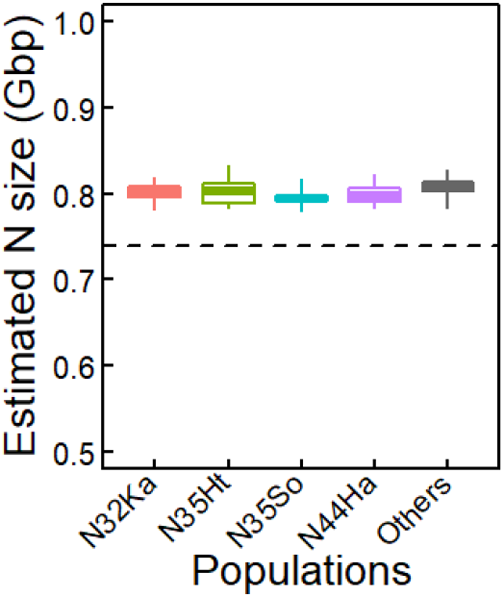
Similar genome sizes among 72 strains in different populations. Box plot displays a median line, interquartile range boxes, min/max whiskers of estimated genome sizes of the strains in a population (N32Ka, N35Ht, N35So, and N44Ha). “Others” includes 16 populations, which are indicated by triangles in Figure 1A. The genome size of a strain was estimated by flow cytometer. A dashed line represents the estimated genome size of *L. aequinoctialis* 6746 strain^33^ isolated in California, USA. No significant difference was detected based on pairwise Wilcoxon test.

**Figure S3.**
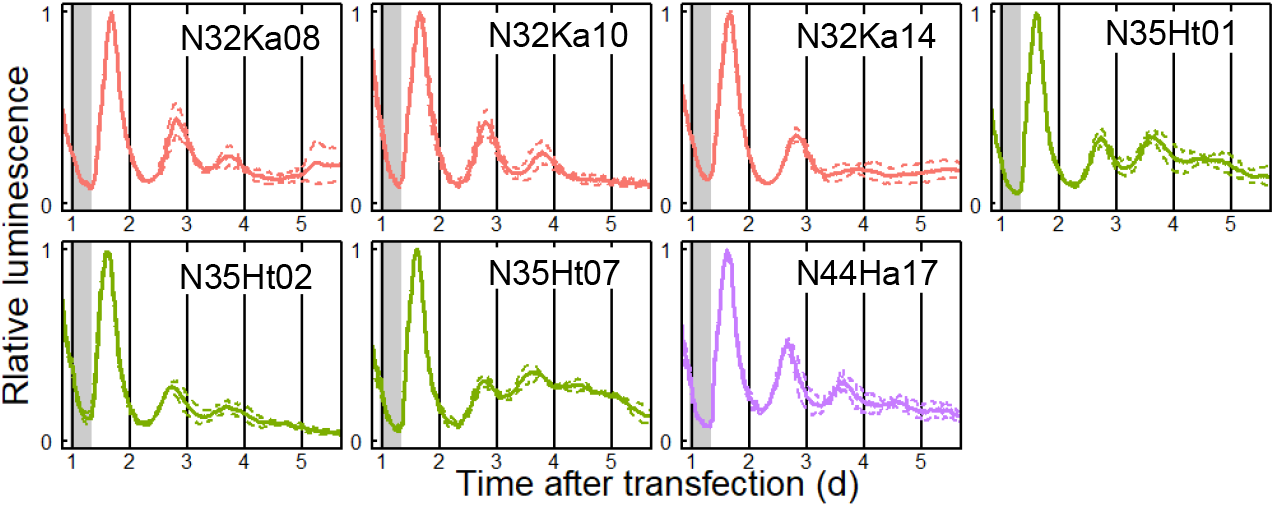
Luminescence rhythms of the seven strains excluded from the analysis. Luminescence intensity at each time point was normalized by the maximum intensity (first peak value). Solid and dashed lines represent mean and SD of three replications, respectively. Same populations are represented by the same colors.

**Figure S4.**
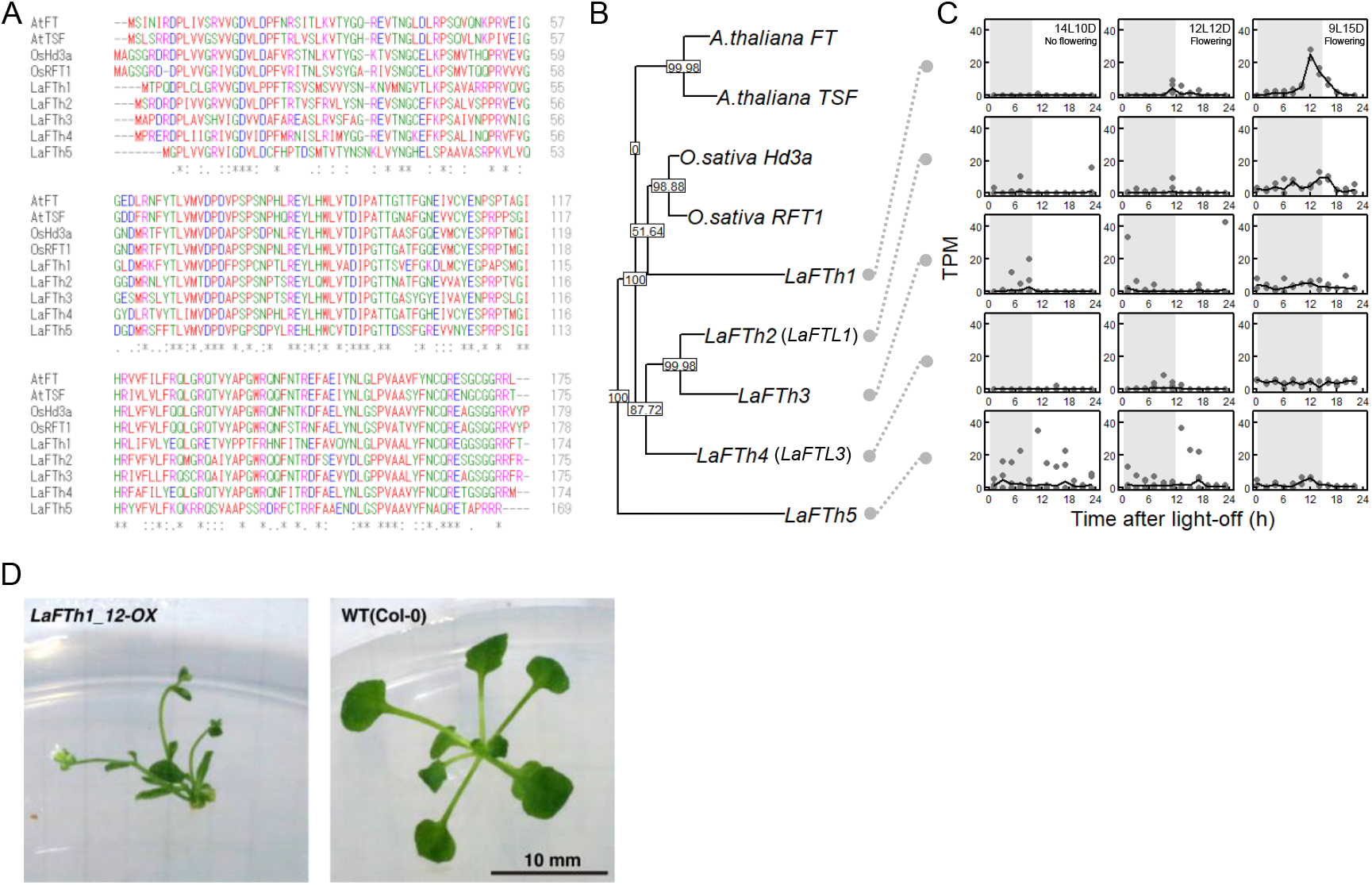
FT homologues of *L. aequinoctialis* Nd strain. **A,** Multiple amino acid sequence alignment of the five FT homologues (LaFTh1-5) with the florigen of *Arabidopsis thaliana* (AtFT and AtTSF) and *Oryza sativa* (OsHd3a and OsRFT1). Colors represent physicochemical properties of residues. **B,** Maximum likelihood phylogenetic tree constructed with PhyML algorithm using full-length amino acid sequences. **C,** The expression patterns of five FT homologues (LaFTh1 to LaFTh5 from top to bottom) under three photoperiods (14L10D, 12L12D, 9L15D from right to left). The transcripts per kilobase million (TPM) values are plotted. Two (9L15D) or four (12L12D and 14L10D) experiments were performed for each photoperiodic condition. Black lines represent the mean values at each time point. Gray boxes indicate dark period. **D**, Early flowering phenotype of *Arabidopsis thaliana* carrying an *LaFTh1* overexpression construct. Plants of an *LaFTh1* overexpression line (T2 generation) (left) and Col-0 wild type (right) on an agar plate at 22 °C and 24 days (10L14D) after sowing.

**Table S1.** Summary of twenty sampling sites.

| # | Population name | Symbol in Figs | Sampling site | Water system | Latitude | Longitude | Number of isolated strains |
| --- | --- | --- | --- | --- | --- | --- | --- |
| 1 | N44Ha | N44Ha | Asahikawa, Hokkaido | Paddy field | 43.80468 | 142.45088 | 12 |
| 2 | N43Hi | Others North | Ishikari, Hokkaido | Paddy field | 43.1943 | 141.45465 | 2 |
| 3 | N41Ac | Others North | Hirosaki castle, Aomori | Castle moat | 40.60337 | 140.46717 | 2 |
| 4 | N40Im | Others North | Miyako, Iwate | Paddy field | 39.56776 | 141.93836 | 1 |
| 5 | N36Ky | Others Middle | Yokohama, Kanagawa | Paddy field | 35.52783 | 139.52045 | 1 |
| 6 | N35At | Others Middle | Togo, Aichi | Paddy field | 35.11062 | 137.083 | 1 |
| 7 | N35Ki | Others Middle | Iwakura, Kyoto | Paddy field | 35.07706 | 135.78781 | 2 |
| 8 | N35Ht | N35Ht | Taka, Hyogo | Paddy field | 35.07304 | 134.90231 | 11 |
| 9 | N35Kk | Others Middle | Kyoto, Kyoto | Paddy field | 35.07206 | 135.74302 | 1 |
| 10 | N35Sb | Others Middle | Biwako, Shiga | Lake | 35.04908 | 135.87446 | 3 |
| 11 | N35Kn | Others Middle | Nougaku, Kyoto | Paddy field | 35.03259 | 135.7836 | 3 |
| 12 | N35Kr | Others Middle | Kamogawa river, Kyoto | Rivier | 34.9896 | 135.76769 | 1 |
| 13 | N35So | N35So | Otsu, Shiga | Paddy field | 34.95615 | 135.95493 | 11 |
| 14 | N35Oi | Others Middle | Ibaraki, Osaka | Paddy field | 34.85884 | 135.56461 | 3 |
| 15 | N35Nn | Others Middle | NAIST, Nara | Paddy field | 34.7317 | 135.73142 | 2 |
| 16 | N35Nk | Others Middle | Kizugawa, Nara | Paddy field | 34.7242 | 135.8158 | 1 |
| 17 | N35Si | Others Middle | Iwata, Shizuoka | Paddy field | 34.72128 | 137.89284 | 1 |
| 18 | N34Mi | Others Middle | Ise, Mie | Paddy field | 34.49805 | 136.67711 | 1 |
| 19 | N32Ka | N32Ka | Aira, Kagoshima | Paddy field | 31.77734 | 130.56278 | 12 |
| 20 | N32Mk | Others South | Kushimoto, Miyazaki | Paddy field | 31.51068 | 131.25419 | 1 |

## Methods

### Plant materials and growth conditions

Seventy-two *L. aequinoctialis* strains were collected from 20 sites in Japan (Table S1). The collected plants were sterilized by washing with 5% sodium hypochlorite for several minutes, followed by washing with sterilized water. From the successfully sterilized plants, a single colony was isolated. Only one strain was isolated from a paddy field, and a maximum of 12 paddy fields were selected at each site. These strains were maintained aseptically in a growth medium (NF medium containing 1% sucrose^33^) under long-day conditions (15L9D; approximately 50 μmol m^-2^ s^-1^) in an incubator (LH350-SP, NK system, Japan) at 25 ± 1 °C. *L. aequinoctialis* 6746 ^33^ and *Lemna gibba* p8L ^33^ strains were maintained in the same conditions. Before the experiments, the plants were grown in a flask or 6-well plastic plate filled with 30 mL or 8 mL of growth medium, respectively.

### Genome size estimation

The relative genome sizes of *L. aequinoctialis* strains were estimated based on 4’,6-diamidino-2-phenylindole diacetate dye flow cytometry (DAPI-FCM) using a Partec CyStain UV Precise P reagent kit (Sysmex Partec GmbH, Germany) and a flow cytometer (Partec Ploidy Analyzer PA-I, Sysmex Partec GmbH). *Lemna gibba* p8L^33^ was used as the standard. The relative genome sizes of th*e L. aequinoctialis* strains were calculated as the relative peak positions. The genome size of *L. gibba* has been estimated to be 486 Mbp.^43^

### Frond-length measurements

The top view image of the plants was captured using a digital camera (EOS 5D mark3 [Canon, Japan] with SP 90 mm Di MACRO [Tamron, Japan] or EOS Kiss X5 with EF-S55-250 mm [Canon]). The frond length was manually measured as the length from the base to the apex of the mother frond using ImageJ 1.53c software (Figure S1A).^51^ The frond lengths of the eight colonies were measured for each strain.

### Measurement of the critical day length in photoperiodic flowering

A colony consisting of three or four visible fronds was placed in each well of a 24-well plastic plate filled with 2 mL of growth medium. These plants were subjected to a one-week photoperiodic treatment with day lengths of 9, 11, 11.5, 12, 12.5, 13, 13.5, 14, 14.5, and 15 h in an eight-chamber incubator (LH30-8-CT, NK system, Japan). The temperature was maintained at 25 ± 1 °C and the light intensity was approximately 30 μmol m^-2^ s^-1^. After treatment, the plants were bleached with 70% ethanol. The number of flowering and non-flowering fronds was counted for each colony under a stereoscopic microscope. Flowering rate was calculated as the percentage of flowering fronds to the total number of fronds in at least two replicates. The critical day length was determined as the day length where 50% of the maximum flowering rate (Fmax) was expected in a piecewise linear function for the obtained flowering rate (Figure 1D).

### Meteorological data

Temperature data at each site were obtained from the Automated Meteorological Data Acquisition System (AMeDAS) data download service (https://www.data.jma.go.jp/risk/obsdl/index.php). Day lengths were calculated using the online service of the National Astronomical Observatory of Japan (https://eco.mtk.nao.ac.jp/cgi-bin/koyomi/koyomix_en.cgi).

### Monitoring luminescence rhythms

Luminescence monitoring using a circadian luminescent reporter was performed as previously described^33^. Plasmid DNA carrying the luciferase reporter gene *pUC-AtCCA1::LUC*+ (*AtCCA1::LUC*^34^) was coated on 0.48 mg of gold particles (1.0 mm diameter, Bio-Rad) and introduced into plants laid on a 60-mm plastic dish using a particle bombardment system (PDS-1000/He, Bio-Rad) according to the manufacturer’s instructions (vacuum, 27 mmHg; helium pressure, 450 psi). After particle bombardment, the plants were divided into three 35-mm dishes (approximately four colonies per dish) filled with 4 mL of growth medium containing D-luciferin (0.2 mM potassium salt, Wako). A luminescence dish-monitoring system with photomultiplier tubes (H7360-01; Hamamatsu Photonics K.K., Japan) was used for the luminescence measurements. To reduce the background chlorophyll fluorescence, a short-pass filter (SVO630; Asahi Spectra, Japan) was set at the detection site of the photomultiplier tubes. Each dish was subjected to 30 s of measurements every 20 min. The monitoring system was placed in an incubator (KCLP-1000I-CT; NK System, Japan) with fluorescent lamps (FL20SSW/18; Mitsubishi Electric Co., Japan). The temperature was maintained at 25 ± 1 °C and the light intensity was approximately 30 μmol m^-2^ s^-1^.

### Time-series analysis

Peak detection and period estimation were performed as previously described.^33, 52^ The peak positions were estimated by local quadratic curve fitting. For period estimation, the obtained luminescence time series were detrended by subtracting the 24-h moving average. Thereafter, the amplitude was normalized by dividing by the 24-h moving standard deviation. The normalized time series of 60 to 132 h was analyzed using fast Fourier transform-nonlinear least squares (FFT-NLLS) to determine the period.^35^ FFT-NLLS was based on a multicomponent cosine fit. Rhythm significance was estimated by a relative amplitude error (RAE) that increases from 0 to 1 as the rhythm approached statistical insignificance. These analyses were performed using R scripts developed and run with R 3.6.3 (http://r-project.org/).^53^

### RNA-seq analysis

Th*e L. aequinoctialis* Nd strain (6^th^ generation of self-fertilization of the N35Kn04 strain)^33^ was used for RNA-seq analysis. Plants were maintained under 15L9D conditions (approximately 50 μmol m^-2^ s^-1^, white LED T5LT20W, Beamtec, Japan) in an incubator (HCLP-880, NK system, Japan) at 25 ± 1 °C. For the 14L10D and 12L12D experiments, the plants were divided into 48 x 35-mm dishes (four colonies per dish) with 4 mL growth medium per dish. After one week of photoperiodic treatment (14L10D or 12L12D), plants in each dish were collected every 2 h for 24 h (four replicates per sampling time). For the 9L15D experiment, plants were pre-cultured under 9L15D conditions for one week. Thereafter, these plants were divided into 24 × 35-mm dishes (six colonies per dish) with 4 mL of growth medium per dish. After one week of photoperiodic treatment (9L15D), plants in each dish were collected every 2 h for 24 h (two replicates at each sampling time). The collected plants were wiped of moisture with paper, then immediately immersed in RNAlater solution (Sigma Aldrich) at each time point and stored at −20 °C before RNA extraction. RNA-seq library preparation was performed based on the BrAD-seq protocol, as follows.^54^ Plants were homogenized with a lysate buffer using a multi-bead shocker (MB755U, Yasui Kikai, Japan). mRNA was extracted from the lysate using magnetic streptavidin beads (New England Biolabs) with a biotin-20nt-20T oligo. The mRNA was subjected to heat fragmentation (94 °C for 1.5 min) and converted to cDNA using RevertAid Reverse Transcriptase (Thermo Fisher) with a 3-prime priming adapter with a random octamer sequence. The 5-prime adapter was then added to the cDNA and captured by the terminus of the RNA-cDNA hybrid. Its incorporation into a complete library molecule was catalyzed by DNA polymerase I (Thermo Fisher). The library was prepared using 16 (9L15D samples) or 18 (14L10D and 12L12D samples) PCR cycles with KAPA HiFi HS ReadyMix (Kapa Biosystems). The fragment size was selected to range from 300 bp to 600 bp using Ampure magnetic beads (Beckman Coulter). The quality of the library was checked using a bioanalyzer (Agilent). Paired-end 150 bp sequencing was conducted using the HiSeq system (Illumina). The obtained sequences (total 554 million reads for the 120 samples) were filtered using Trimmomatic v0.39 with option “LEADING:24 TRAILING:24 SLIDINGWINDOW:30:20 AVGQUAL:20 MINLEN:100”.^55^ The filtered sequences were assembled *de novo* into 322,899 contigs (isoforms of 186,102 genes) using Trinity v2.8.4.^56^ The 183,102 contigs (isoforms of 84,206 genes) that were longer than 400 bp were used as a reference sequence for mapping and read counting using RSEM v1.3.1 with bowtie2 v2.4.2.^57, 58^ The mapped reads were 3.46 ± 1.22 million reads / sample (mean ± SD).

### Detection of *FT* homologues

*FT* homologues in the reference sequence (183,102 contigs) were searched using tblastn 2.11.0+^59^ with a query for the OsHd3a amino acid sequence (UniProt-ID: Q93WI9). The top five *FT* homologues were named as *LaFTh1-5* based on their position in the maximum likelihood phylogenetic tree with their estimated amino acid sequences and AtFT (UniProt-ID: Q9SXZ2), AtTSF (UniProt-ID: Q9S7R5), OsHd3a (UniProt-ID: Q93WI9), and OsRFT1(UniProt-ID: Q8VWH2) (Figure S4B). LaFTh2 and LaFTh4 have the same sequence as previously reported FT homologues, LaFTL1 and LaFTL3, respectively.^60^ A phylogenetic tree was constructed using the online tool ClustalW (https://www.genome.jp/tools-bin/clustalw) using the PhyML algorithm. Colored multiple amino acid sequence alignments were generated using the online tool Clustal Omega (https://www.ebi.ac.uk/Tools/msa/clustalo/).

### qPCR analysis

Eight to 12 colonies grown under 15L9D conditions were placed in a 35-mm dish with 4 mL of growth medium and maintained under the same conditions for a day, after which they were released to constant darkness at the end of 15 h of light and sampled every hour from 9 to 15 h after the light-off. The collected plants were wiped of moisture with paper, then immediately immersed in RNAlater solution (Sigma Aldrich) at each time point and stored at −20 °C before RNA extraction. Total RNA was extracted using the NucleoSpin RNA Plant (MACHEREY-NAGEL GmbH & Co.) and converted to cDNA using ReverTra Ace qPCR RT Master Mix (TOYOBO). qPCR was performed using the StepOnePlus Real-Time PCR System (Applied Biosystems) with THUNDERBIRD SYBR qPCR Mix (TOYOBO). An *ACT2* homologue in *de novo* assembled sequences was determined using tblastn 2.11.0+ with a query of the OsACT2 amino acid sequence (UniProt-ID: A3C6D7) and used as a reference gene for the delta-delta Ct method. Primers for qPCR analysis were designed according to the *de novo* assembled sequences as follows: LaFTh1_fwd, 5’-ACCCTACCCTTAGAGAATATCTGC-3’; LaFTh1_rev, 5’ -TAGGTGCTGGGCCTTCATAG-3’, LaACT2_fwd, 5’-ACACAGTGCCCATCTATGAAGG-3’; LaACT2_rev, 5’-AGTAGCCTCGTTCGGTTAGGATC-3’.

### *LaFTh1* overexpression in *Arabidopsis thaliana*

To generate *LaFTh1* overexpressing transgenic *Arabidopsis* plants, the coding region of *LaFTh1* was amplified from cDNA derived from *L. aequinoctialis* Nd using the following primers: 5’-CGCGGCCGCCACCATGACACCTCAGGATCCCTTGTG-3’, and 5’-CGGCGCGCCCTAGGTAAACCGCCTTCCACCAG-3’. The amplified PCR fragment was digested with *NotI* and *AscI* restriction enzymes and integrated into the pENTR-MCS cloning vector [pENTR-D/TOPO vector (Invitrogen) containing the pBluescript II multi-cloning sites] using the conventional linker ligation method. The sequence of *LaFTh1* CDS was transferred into the pK7WG2 overexpression binary vector,^61^ which contained the *CaMV35S* promoter-driven expression cassette, using Gateway LR clonase II (Invitrogen). This construct was introduced into *Arabidopsis* wild-type plants Col-0 by the *Agrobacterium tumefaciens-mediated* floral dipping method.^62^ Primary transformants (T1) were selected on 1x Murashige and Skoog culture media containing 1% sucrose, 0.8% agar, and 25 μg mL^-1^ kanamycin sulfate. Seeds were sterilized using chlorine gas, stratified for 3 days at 4 °C, and then grown under short-day (10L14D) conditions to analyze the flowering phenotype in parallel with the selection process. Seedlings developing relatively long primary and lateral roots were regarded as kanamycin-resistant transgenic plants and transferred to a 1:1 mixture of vermiculite and soil (Nihon Hiryo Co., Ltd.). Several independent lines (*N* > 10) were isolated for further analysis of their morphological and flowering phenotypes. Most primary transformants could not produce mature siliques or seeds; however, some transgenic plants produced T2 seeds. Flowering phenotypes were also observed in the T2 generation.

### Statistics

All boxplots in this manuscript display median lines, interquartile range boxes, and min/max whiskers. Different letters indicate significant differences based on the pairwise Wilcoxon test (*p* < 0.05). The pairwise Wilcoxon test was performed using the R built-in function “pairwise.wilcox.test” with *P* value adjustment using the Holm method. In correlation analysis, the Pearson’s correlation coefficient and *P* value was calculated by the R-built-in function “cor.test”.

## References

1. Barrett, R.D.H., and Schluter, D. (2008). Adaptation from standing genetic variation. Trends Ecol. Evol. 23, 38–44.

2. Alonso-Blanco, C., Aarts, M.G.M., Bentsink, L., Keurentjes, J.J.B., Reymond, M., Vreugdenhil, D., and Koornneef, M. (2009). What has natural variation taught us about plant development, physiology, and adaptation? Plant Cell 21, 1877–1896.

3. Matuszewski, S., Hermisson, J., and Kopp, M. (2015). Catch me if you can: adaptation from standing genetic variation to a moving phenotypic optimum. Genetics 200, 1255–1274.

4. Boyle, E.A., Li, Y.I., and Pritchard, J.K. (2017). An Expanded view of complex traits: From polygenic to omnigenic. Cell 169, 1177–1186.

5. Sella, G., and Barton, N.H. (2019). Thinking about the evolution of complex traits in the era of genome-wide association studies. Annu. Rev. Genomics Hum. Genet. 20, 461–493.

6. Feiner, N., Brun-Usan, M., and Uller, T. (2021). Evolvability and evolutionary rescue. Evol. Dev. 23, 308–319.

7. Helm, B., Visser, M.E., Schwartz, W., Kronfeld-Schor, N., Gerkema, M., Piersma, T., and Bloch, G. (2017). Two sides of a coin: ecological and chronobiological perspectives of timing in the wild. Philos. Trans. R. Soc. Lond. B. Biol. Sci. 372, 20160246.

8. Michael, T.P., Park, S., Kim, T.S., Booth, J., Byer, A., Sun, Q., Chory, J., and Lee, K. (2007). Simple sequence repeats provide a substrate for phenotypic variation in the *Neurospora crassa* circadian clock. PloS One 2, e795.

9. Brown, S.A., Kunz, D., Dumas, A., Westermark, P.O., Vanselow, K., Tilmann-Wahnschaffe, A., Herzel, H., and Kramer, A. (2008). Molecular insights into human daily behavior. Proc. Natl. Acad. Sci. U. S. A. 105, 1602–1607.

10. Pivarciova, L., Vaneckova, H., Provaznik, J., Wu, B.C.H., Pivarci, M., Peckova, O., Bazalova, O., Cada, S., Kment, P., Kotwica-Rolinska, J., et al. (2016). Unexpected Geographic Variability of the Free Running Period in the Linden Bug *Pyrrhocoris apterus*. J. Biol. Rhythms 31, 568–576.

11. Salmela, M.J., and Weinig, C. (2019). The fitness benefits of genetic variation in circadian clock regulation. Curr. Opin. Plant Biol. 49, 86–93.

12. Hut, R.A., Paolucci, S., Dor, R., Kyriacou, C.P., and Daan, S. (2013). Latitudinal clines: An evolutionary view on biological rhythms. Proc. R. Soc. B Biol. Sci. 280, 1–9.

13. Kubota, A., Shim, J.S., and Imaizumi, T. (2015). Natural variation in transcriptional rhythms modulates photoperiodic responses. Trends Plant Sci. 20, 259–261.

14. Yukawa, I., and Takimoto, A. (1976). Flowering response of *Lemna paucicostata* in Japan. Bot. Mag. Tokyo 89, 241–250.

15. Pittendrigh, C.S. (1960). Circadian rhythms and the circadian organization of living systems. Cold Spring Harb. Symp. Quant. Biol. 25, 159–184.

16. Saini, R., Jaskolski, M., and Davis, S.J. (2019). Circadian oscillator proteins across the kingdoms of life: Structural aspects. BMC Biol. 17, 1–39.

17. Nagel, D.H., and Kay, S.A. (2012). Complexity in the wiring and regulation of plant circadian networks. Curr. Biol. 22, R648–57.

18. Johnson, C.H., Elliott, J.A., and Foster, R. (2003). Entrainment of circadian programs. Chronobiol. Int. 20, 741–774.

19. Schmal, C., Herzel, H., and Myung, J. (2020). Clocks in the wild: Entrainment to natural light. Front. Physiol. 11, 1–12.

20. Aschoff, J., and Pohl, H. (1978). Phase relations between a circadian rhythm and its zeitgeber within the range of entrainment. Naturwissenschaften 65, 80–84.

21. Lankinen, P. (1986). Geographical variation in circadian eclosion rhythm and photoperiodic adult diapause in *Drosophila littoralis*. J. Comp. Physiol. A 159, 123–142.

22. Graf, A., Schlereth, A., Stitt, M., and Smith, A.M. (2010). Circadian control of carbohydrate availability for growth in *Arabidopsis* plants at night. Proc. Natl. Acad. Sci. U. S. A. 107, 9458–9463.

23. Rémi, J., Merrow, M., and Roenneberg, T. (2010). A circadian surface of entrainment: varying T, τ, and photoperiod in *Neurospora crassa*. J. Biol. Rhythms 25, 318–328.

24. Granada, A.E., Bordyugov, G., Kramer, A., and Herzel, H. (2013). Human chronotypes from a theoretical perspective. PloS One 8, e59464.

25. Dominoni, D.M., Helm, B., Lehmann, M., Dowse, H.B., and Partecke, J. (2013). Clocks for the city: circadian differences between forest and city songbirds. Proceedings. Biol. Sci. 280, 20130593.

26. Jiménez-Gómez, J.M., Lin, T., Srinivasan, A., Maloof, J.N., Müller, N.A., Ranjan, A., Ryngajllo, M., Sinha, N.R., Huang, S., West, D., et al. (2015). Domestication selected for deceleration of the circadian clock in cultivated tomato. Nat. Genet. 48, 89–93.

27. Ray, P.M., and Alexander, W.E. (1966). Photoperiodic adaptation to latitude in *Xanthium strumarium*. Am. J. Bot. 53, 806–816.

28. Imamura, S., Muramatsu, M., Kitajo, S.I., and Takimoto, A. (1966). Varietal difference in photoperiodic *Pharbitis nil*. Dot. Mag. Tokyo 79, 714–721.

29. Katayama, T. (1977). Studies on the photoperiodism in the genus *Oryza*. Japan Agric. Res. Q. 11, 12–17.

30. Natuhara, Y. (2013). Ecosystem services by paddy fields as substitutes of natural wetlands in Japan. Ecol. Eng. 56, 97–106.

31. Fujita, G., Naoe, S., and Miyashita, T. (2015). Modernization of drainage systems decreases gray-faced buzzard occurrence by reducing frog densities in paddy-dominated landscapes. Landsc. Ecol. Eng. 11, 189–198.

32. Landolt, E. (1986). Biosystematic investigation in the family of duckweeds (“*Lemnaceae*”). Vol. 2: the family of *“Lemnaceae”* – a monographic study. Volume 1 Veröffentlichungen des Geobotanischen Institutes der ETH, Stiftung Rubel, Zürich 78, 142–146.

33. Muranaka, T., Okada, M., Yomo, J., Kubota, S., and Oyama, T. (2015). Characterisation of circadian rhythms of various duckweeds. Plant Biol. 17, 66–74.

34. Nakamichi, N., Ito, S., Oyama, T., Yamashino, T., Kondo, T., and Mizuno, T. (2004). Characterization of plant circadian rhythms by employing *Arabidopsis* cultured cells with bioluminescence reporters. Plant Cell Physiol. 45, 57–67.

35. Zielinski, T., Moore, A.M., Troup, E., Halliday, K.J., and Millar, A.J. (2014). Strengths and limitations of period estimation methods for circadian data. PloS One 9, e96462.

36. Hayama, R., Agashe, B., Luley, E., King, R., and Coupland, G. (2007). A circadian rhythm set by dusk determines the expression of FT homologs and the short-day photoperiodic flowering response in Pharbitis. Plant Cell 19, 2988–3000.

37. Hut, R.A., and Beersma, D.G.M. (2011). Evolution of time-keeping mechanisms: Early emergence and adaptation to photoperiod. Philos. Trans. R. Soc. B Biol. Sci. 366, 2141–2154.

38. Katayama, N., Baba, Y.G., Kusumoto, Y., and Tanaka, K. (2015). A review of post-war changes in rice farming and biodiversity in Japan. Agric. Syst. 132, 73–84.

39. Michael, T.P., Salomé, P.A., Yu, H.J., Spencer, T.R., Sharp, E.L., McPeek, M.A., Alonso, J.M., Ecker, J.R., and McClung, C.R. (2003). Enhanced fitness conferred by naturally occurring variation in the circadian clock. Science. 302, 1049–1053.

40. Rieseberg, L.H., Widmer, A., Arntz, A.M., Burke, J.M., Carr, D.E., Abbott, R.J., and Meagher, T.R. (2003). The genetic architecture necessary for transgressive segregation is common in both natural and domesticated populations. Philos. Trans. R. Soc. B Biol. Sci. 358, 1141–1147.

41. Hamann, E., Pauli, C.S., Joly-Lopez, Z., Groen, S.C., Rest, J.S., Kane, N.C., Purugganan, M.D., and Franks, S.J. (2021). Rapid evolutionary changes in gene expression in response to climate fluctuations. Mol. Ecol. 30, 193–206.

42. Beppu, T., and Takimoto, A. (1981). Geographical distribution and cytological variation of *Lemna paucicostata* Hegelm. In Japan. Bot. Mag. Tokyo 94, 11–20.

43. Wang, W., Kerstetter, R.A., and Michael, T.P. (2011). Evolution of genome size in duckweeds *(Lemnaceae)*. J. Bot. 2011, 1–9.

44. Sémon, M., and Wolfe, K.H. (2007). Consequences of genome duplication. Curr. Opin. Genet. Dev. 17, 505–512.

45. Selmecki, A.M., Maruvka, Y.E., Richmond, P.A., Guillet, M., Shoresh, N., Sorenson, A.L., De, S., Kishony, R., Michor, F., Dowell, R., et al. (2015). Polyploidy can drive rapid adaptation in yeast. Nature 519, 349–351.

46. Greenham, K., and McClung, C.R. (2015). Integrating circadian dynamics with physiological processes in plants. Nat. Rev. Genet. 16, 598–610.

47. Urbanová, V., Bazalová, O., Vaněčková, H., and Dolezel, D. (2016). Photoperiod regulates growth of male accessory glands through juvenile hormone signaling in the linden bug, *Pyrrhocoris apterus*. Insect Biochem. Mol. Biol. 70, 184–190.

48. Dixon, L.E., Hodge, S.K., van Ooijen, G., Troein, C., Akman, O.E., and Millar, A.J. (2014). Light and circadian regulation of clock components aids flexible responses to environmental signals. New Phytol. 203, 568–577.

49. Nagel, D.H., and Kay, S.A. (2012). Complexity in the wiring and regulation of plant circadian networks. Curr. Biol. 22, R648–57.

50. Sun, L., Dong, A., Griffin, C., and Wu, R. (2021). Statistical mechanics of clock gene networks underlying circadian rhythms. Appl. Phys. Rev. 8, 021313.

51. Schneider, C.A., Rasband, W.S., and Eliceiri, K.W. (2012). NIH Image to ImageJ: 25 years of image analysis. Nat. Methods 9, 671–675.

52. Muranaka, T., and Oyama, T. (2016). Heterogeneity of cellular circadian clocks in intact plants and its correction under light-dark cycles. Sci. Adv. 2, e1600500.

53. Ihaka, R., and Gentleman, R. (1996). R: A language for data analysis and graphics. J. Comput. Graph. Stat. 5, 299–314.

54. Townsley, B.T., Covington, M.F., Ichihashi, Y., Zumstein, K., and Sinha, N.R. (2015). BrAD-seq: Breath Adapter Directional sequencing: A streamlined, ultra-simple and fast library preparation protocol for strand specific mRNA library construction. Front. Plant Sci. 6, 1–11.

55. Bolger, A.M., Lohse, M., and Usadel, B. (2014). Trimmomatic: A flexible trimmer for Illumina sequence data. Bioinformatics 30, 2114–2120.

56. Grabherr, M.G., Haas, B.J., Yassour, M., Levin, J.Z., Thompson, D.A., Amit, I., Adiconis, X., Fan, L., Raychowdhury, R., Zeng, Q., et al. (2011). Full-length transcriptome assembly from RNA-Seq data without a reference genome. Nat. Biotechnol. 29, 644–652.

57. Li, B., and Dewey, C.N. (2011). RSEM: accurate transcript quantification from RNA-Seq data with or without a reference genome. BMC Bioinformatics 12, 323.

58. Langmead, B., and Salzberg, S.L. (2012). Fast gapped-read alignment with Bowtie 2. Nat. Methods 9, 357–359.

59. Altschul, S.F., Gish, W., Miller, W., Myers, E.W., and Lipman, D.J. (1990). Basic local alignment search tool. J. Mol. Biol. 215, 403–410.

60. Yoshida, A., Taoka, K., Hosaka, A., Tanaka, K., Kobayashi, H., Muranaka, T., Toyooka, K., Oyama, T., and Tsuji, H. (2021). Characterization of frond and flower development and identification of FT and FD genes from duckweed *Lemna aequinoctialis* Nd. Front. Plant Sci. 12, 697206.

61. Karimi, M., Inzé, D., and Depicker, A. (2002). GATEWAY vectors for *Agrobacterium*-mediated plant.pdf. Trends Plant Sci. 7, 193–195.

62. Zhang, X., Henriques, R., Lin, S.S., Niu, Q.W., and Chua, N.H. (2006). *Agrobacterium*-mediated transformation of *Arabidopsis thaliana* using the floral dip method. Nat. Protoc. 1, 641–646.

